# Dynamic Pocketome of Trace Amine-Associated Receptors

**DOI:** 10.64898/2026.09.14.751399

**Authors:** Clarissa Rienaecker, Alessandro Nicoli, Jana Selent, Antonella Di Pizio

**Affiliations:** Technical University of Munich, Germany; TUM School of Life Sciences, Professorship for Chemoinformatics and Protein Modeling, Lise-Meitner-Str. 34, 85354 Freising, Germany; Leibniz Institute for Food Systems Biology at the Technical University of Munich, Lise-Meitner-Str. 34, 85354 Freising, Germany; Research Programme on Biomedical Informatics (GRIB), Hospital del Mar Medical Research Institute & Pompeu Fabra University, Barcelona, Spain; Atomistic Modeling Center, Munich Data Science Institute, Technical University of Munich, Garching, Germany

**Keywords:** GPCR, molecular dynamics, binding site plasticity, orthosteric pocket, allosteric pocket, transient pocket, chemosensory receptors

## Abstract

Trace amine-associated receptors (TAARs) are class A GPCRs that span two distinct physiological roles: TAAR1 is a CNS drug target, whereas TAAR2 to TAAR9 detect volatile amines in the olfactory epithelium. Recent experimental structures resolve their architecture and ligand-binding mode, but capture only static snapshots, which cannot address how the binding site and the overall pocketome respond to ligand binding. Here, we present a simulation library comprising 26 experimental structures of four human and murine genes in the apo and holo states, each in triplicate (total aggregate time of 156 µs). Cavities were detected and analysed across the entire receptor surface throughout each trajectory. Orthosteric changes did not follow a single direction when comparing the states: apo sites were neither uniformly smaller nor uniformly more flexible than their holo counterparts, instead pointing to a receptor-specific ligand-receptor interplay that propagates beyond the orthosteric pocket. This plasticity is further highlighted by the size composition and stability of allosteric pockets, which proves that some regions are larger in the apo state while others are larger when a ligand is present. Resolving such trends required pockets to be comparable across trajectories, which a novel global identifier (GID) provides. Although agnostic to functional annotation, the GID recovered the orthosteric site as a single region in both states and matched pockets across replicates of the same receptor-state pair. The result is a dynamic pocketome map of the TAAR family obtained by an approach transferable to other membrane proteins.

**Graphical Abstract:** 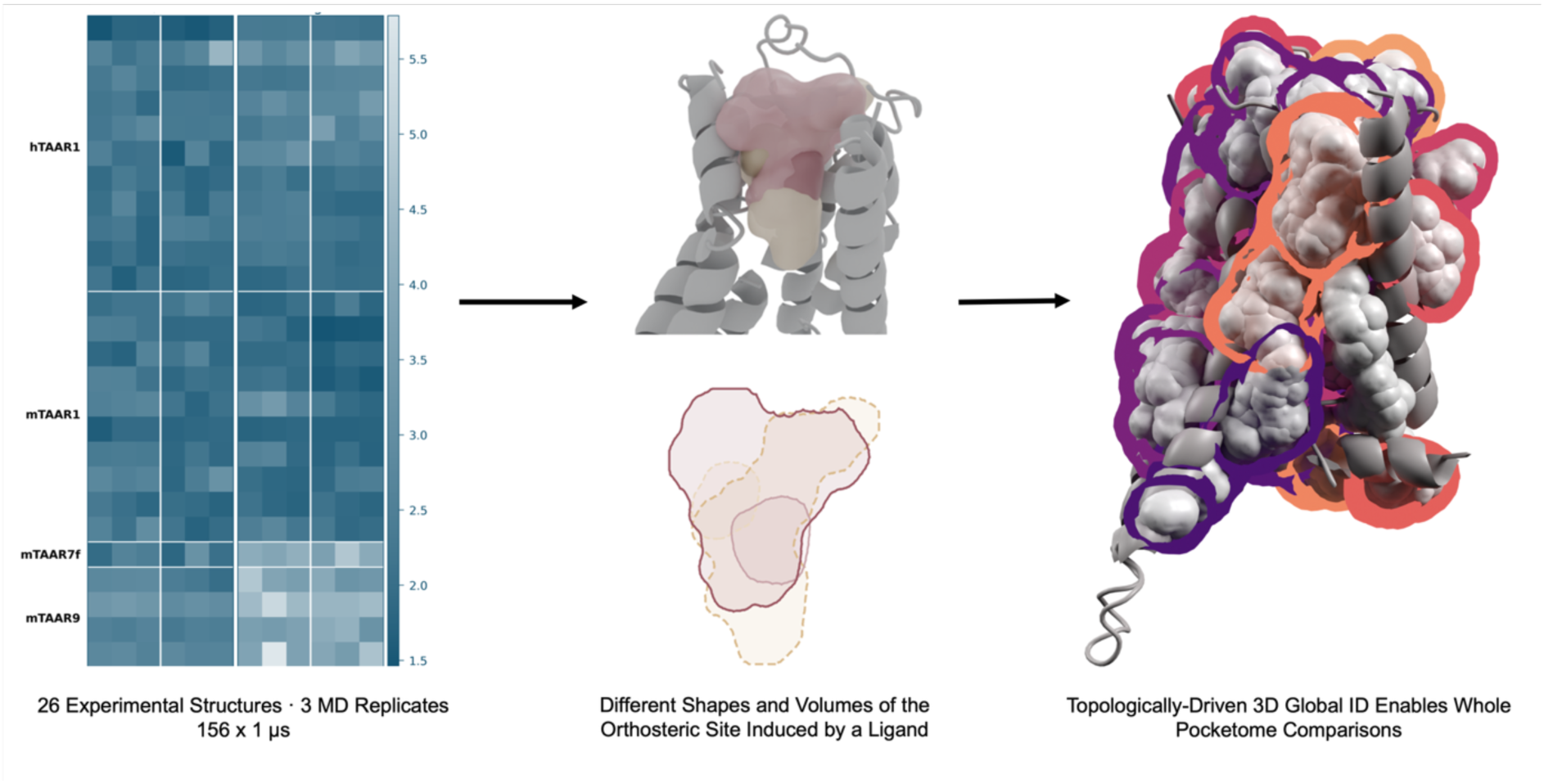

## Introduction

G protein-coupled receptors (GPCRs) serve as the primary interface between the extracellular environment and intracellular effectors, making them crucial to human physiology. With over 800 genomic sequences, GPCRs are the largest family of membrane receptors, with highly diverse architecture and function [1], [2]. GPCRs are grouped into six classes according to two overlapping classification systems. The traditional nomenclature designates these as classes A – F [1], while the GRAFS system, based on phylogenetic analysis and sequence homology, categorises them by their prototypical members: Glutamate, Rhodopsin, Adhesion, Frizzled, and Secretin [1]. While class A (Rhodopsin) is the most extensively studied and serves as the target for 34% of all clinical drugs, a vast majority of the wider GPCR superfamily remains understudied or orphaned [3], [4].

A significant portion of this ‘dark’ GPCR landscape [4] is occupied by the olfactory receptor superfamily, which encompasses over 400 members, with ca. 390 Odorant Receptors (ORs) and six Trace Amine-Associated Receptors (TAARs) [5]. Notably, 88% of ORs remain orphans [6], and their structural architecture diverges significantly from that of canonical class A GPCRs. In contrast, TAARs share greater sequence and structural homology with prototypical class A receptors. TAAR1 is a well-established drug target expressed prominently in the brain [7], [8]. TAAR2 – 9 were historically defined as specialised olfactory sensors, but their expression in extra-nasal tissues suggests a much broader physiological role [9], [10]. This expression profile, coupled with the trial of human TAAR1 agonist ulotaront in clinical trials for the treatment of schizophrenia [11], underscores the significant, yet under-exploited, druggability of the TAAR family. To unlock this therapeutic potential, a detailed and mechanistic understanding of receptor structures and their functional dynamics is required.

Although recent experimental breakthroughs have expanded the pool of available GPCR structures to include the TAAR family [5], these are inherently static snapshots and cannot capture the physiological flexibility required for receptor function [12], [13]. This highlights the need for computational frameworks that can analyse and interpret this data dynamically [14]. To this end, we have developed a comprehensive library of molecular dynamics (MD) simulations to explore the functional conformational space of TAAR structures. By systematically mapping the pockets formed throughout these simulations, we compare these receptors across different genes and species. Furthermore, we introduce a novel global identifier framework to bridge the gap between local, trajectory-specific cavity detection and system-wide structural analysis. By providing a dynamic, high-resolution map of the TAAR pocketome, this study offers a scalable approach for characterising binding sites and facilitates future ligand discovery across this understudied receptor family.

### Results and Discussion

In the following study, we present an extensive MD simulation library, including both apo and holo states across human and murine TAARs. By analysing and characterising the pocketome of each trajectory, we capture the structural plasticity of TAARs. To compare the pocketome across the experiments, we introduce a new global identifier (GID), enabling a structured and automated analysis of conformational changes represented in the pocketome. The new identifier also enables the systematic comparison within genes and between apo and holo states. The complete analysis pipeline is openly available through the Di Pizio Lab GitHub account (https://github.com/DiPizio-Lab/dynamic_pocketome_TAARs) and MD data is accessible through GPCRmd [15] (ID pending) and Zenodo [16] (10.5281/zenodo.22661709 (MD data), 10.5281/zenodo.22709348 (pocketome)).

## 1. TAAR Structural Plasticity

In order to investigate the functional landscape of TAARs, we generated an extensive library of 156 microseconds of MD simulation data. Our dataset covers 26 different TAAR structures from four genes and two species (human and mouse) in both the apo and holo states. With three replicates per condition, this resulted in 156 individual experimental setups (experiment is defined as the unique combination of PDB ID, replicate and state) (Figure 1).

**Figure 1.**
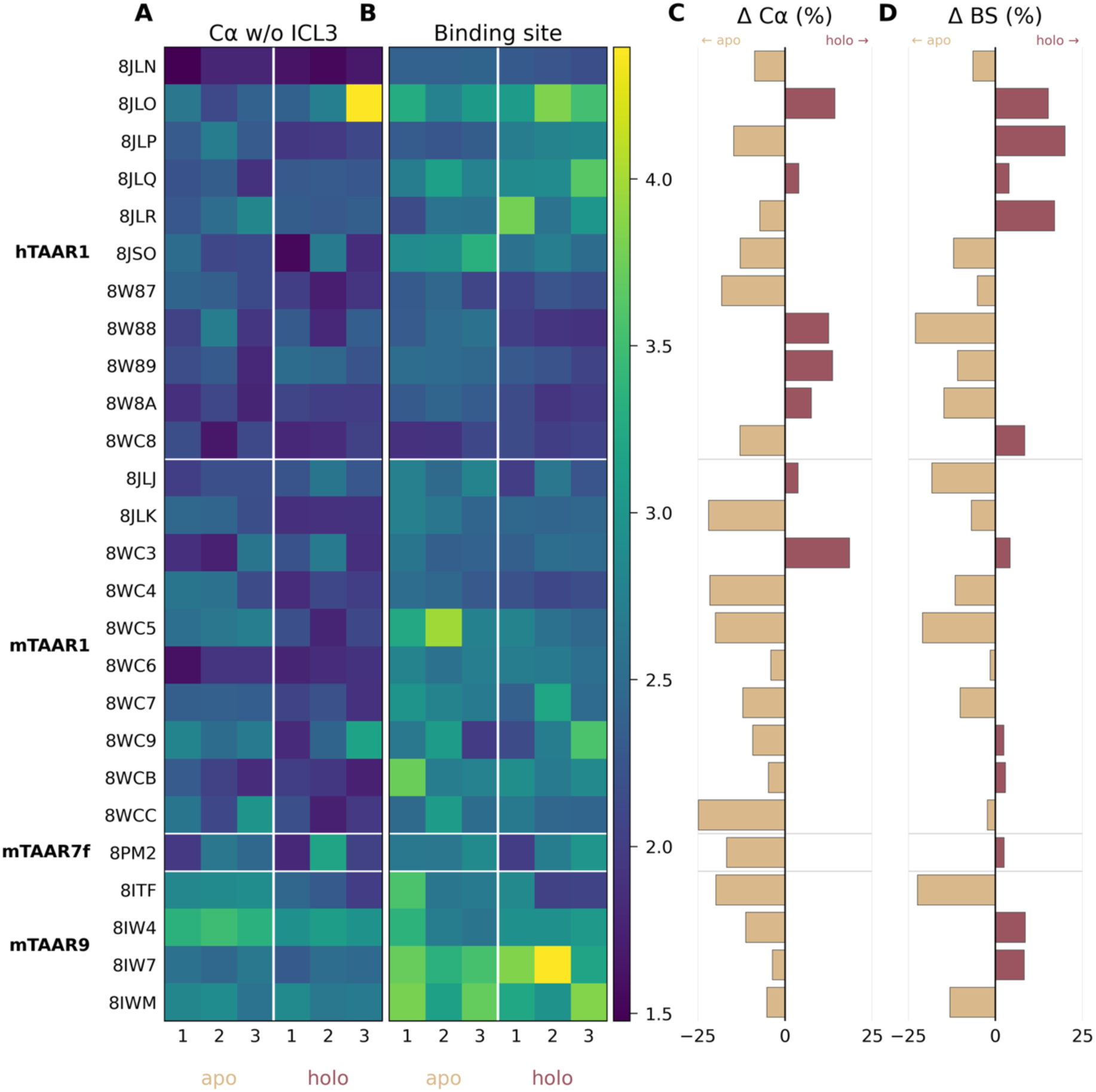
Overview of the simulation workflow and structural analysis of TAARs. A total of 26 PDB structures were simulated in triplicate across two states (apo = without the ligand, holo = with the co-bound ligand), spanning four human and murine genes. (**A**) Heatmap summarising the median RMSD of the Carbon alpha backbone for each replicate. (**B**) Conformational changes of binding site residues in terms of their median RMSD. (**C, D**) Difference between apo and holo RMSD values for the backbone and binding site residues, respectively. All RMSD calculations of the receptor backbone excluded the intracellular loop 3 region (ICL3).

All simulations exhibited stable behaviour across the three replicates, as evidenced by consistent Root Mean Square Deviation (RMSD) profiles (Figure 1A, 1B, Sup Fig 1). A critical observation in our analysis was the significant contribution of intracellular loop 3 (ICL3) to the overall backbone Cα RMSD. Because our simulations were performed in the absence of a G protein, this extended loop remains unconstrained. When excluding the ICL3 region from our calculations, the overall RMSD was reduced by 21% and 20% for apo and holo states, respectively (Sup Fig 1). Of note, one replicate of 8JLO (hTAAR1) shows a higher RMSD than the other simulations, driven by a unique conformational shift in ICL3, which folds back to act like a lid over the intracellular region. Although ICL3 was excluded from calculations, this motion pulls on the otherwise stable receptor during the latter half of the simulation. Since this movement is not seen in any other simulations and would not be possible in the presence of a G protein, this replicate is not representative of the expected physiological ICL3 movement. Nevertheless, it does not affect the following analyses of the helix bundle focused pocketome.

The median RMSD of the binding site residues was higher than the overall backbone Cα RMSD, indicating that local rearrangements in the orthosteric binding site exceed the conformational changes observed for the receptor backbone. It is notable that all murine TAAR9 simulations have a higher median RMSD than most other replicates. This behaviour is mainly driven by the orthosteric binding site residues, as seen in Figure 1B.

To evaluate the impact of ligand binding on receptor dynamics, we compared the RMSD of the binding site side chains between apo and holo states. Ligand occupancy might be expected to restrict the conformational freedom of residues directly interacting with the ligand, consistent with previous observations of ligand-mediated stabilisation of GPCR binding pockets [17]. However, no uniform apo–holo trend was observed across the TAAR structures analysed here (Figure *1*D), nor was consistent correlation with the difference between apo and holo RMSD values for the backbone (Figure *1*C).

While a majority of the analysed structures exhibit higher side-chain mobility in the apo state (negative difference RMSD in the plot, Figure 1D), likely reflecting the lack of steric hindrance from a ligand, a significant subset of receptors shows higher or comparable mobility in the holo state. This variability warrants a careful interpretation of the “apo” state, as these structures were derived by removing the co-bound ligand from the holo-form. Consequently, some apo simulations likely represent a state in which the binding site residues undergo significant conformational relaxation once the ligand constraint is removed. However, the fact that this relaxation is not uniform across all structures — and that some holo simulations remain highly dynamic — suggests that ligand-induced conformational changes are highly dependent on the initial scaffold and the chemical nature of the co-bound ligand. This occurs partly because binding site residues are distributed widely throughout the receptor architecture, including portions of the extracellular loops, allowing the residues to retain local flexibility even when bound.

Furthermore, this observation suggests that ligand binding in TAARs does not merely “lock” the binding pocket in a rigid configuration. Instead, it may induce allosteric rearrangements that propagate dynamics throughout the receptor structure, potentially activating secondary or distal allosteric pockets. Consequently, the variability in our RMSD data underscores that binding site flexibility is highly ligand-dependent and suggests that TAARs may possess a more complex allosteric landscape than previously assumed.

## 2. TAAR Orthosteric Binding site Plasticity

A systematic comparison of the orthosteric binding site across TAARs may provide structural information relevant to receptor de-orphanisation, particularly for poorly characterised family members for which ligand recognition remains unresolved [18]. While RMSD metrics reveal a complex interplay of flexibility across the receptor, they primarily capture atomic-level fluctuations rather than functional cavity dimensions. To better understand how these conformational dynamics influence ligand recognition, we next examined the orthosteric binding site in detail by analysing its volume distribution.

We observed a clear distinction in volume distribution between apo and holo states (Figure 2A). In the majority of our simulations (88%), the median orthosteric pocket volume is significantly larger in the holo state than in the corresponding apo state. This volumetric expansion is further highlighted by pocket ranking: the orthosteric site corresponds to the single largest characterised pocket in 54% of holo simulations, compared to only 19% of apo simulations. In apo state simulations, the binding site can additionally be fragmented into several smaller pockets. This finding aligns with broader observations in GPCR structural biology, where in the absence of a ligand, this cavity frequently “collapses” into a more restricted, low-volume conformation [19]. Notably, only two PDB IDs show a higher volume of their orthosteric site in the apo state than their holo counterparts (Figure 2 B).

**Figure 2.**
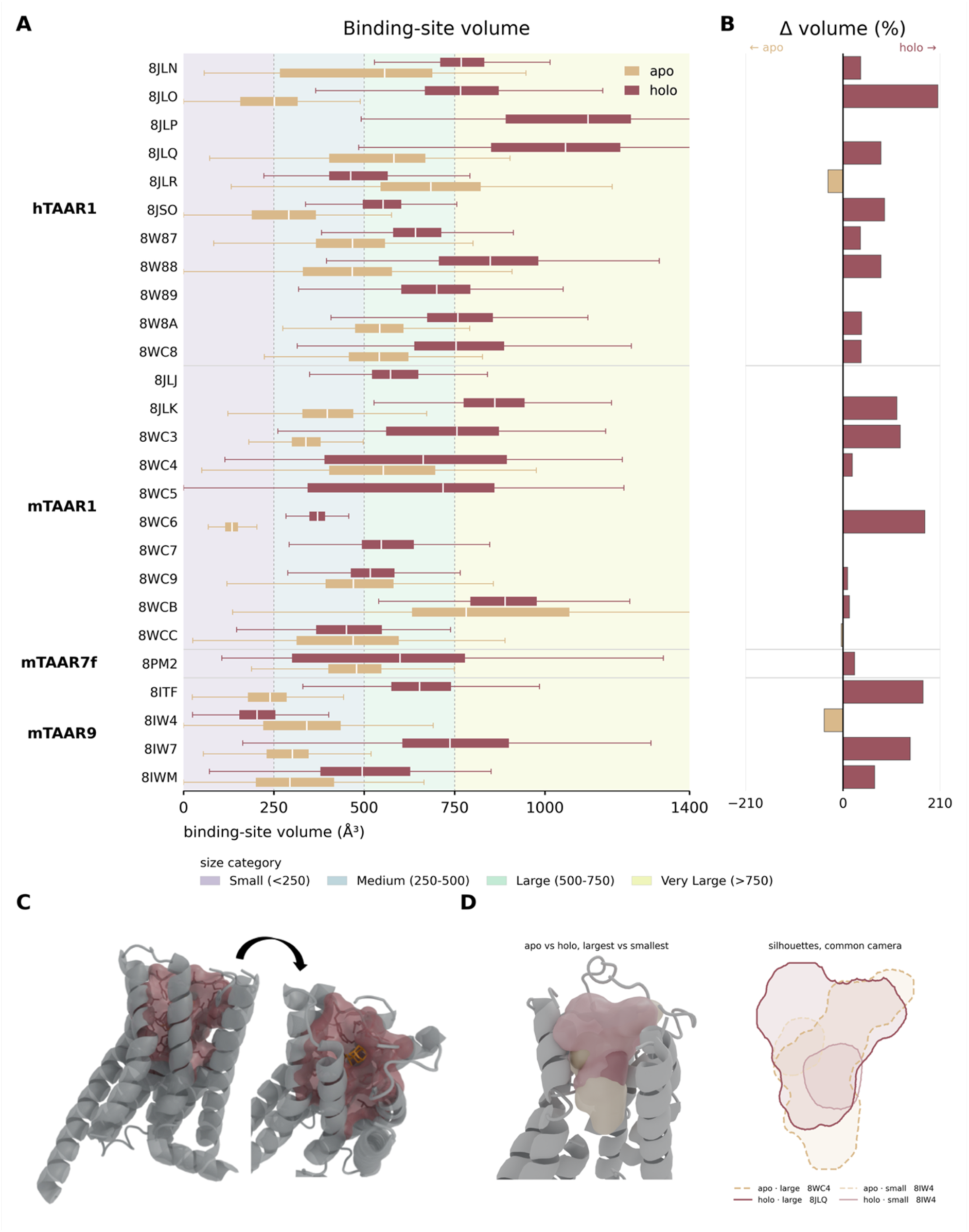
Orthosteric pocket volume dynamics. (**A**) Boxplots of volume distribution over simulation time, including volume categories. (**B**) Differential volume plot highlighting state-specific differences. (**C**) Representative 3D structure of hTAAR1 (8JLR) bound to VRK (compound A77636, depicted in orange) to contextualise the orthosteric site location. (**D**) Structural overlay comparing the extreme (largest and smallest) apo and holo binding site pockets (coloured in sand and mauve, respectively) within a human TAAR1 backbone (with transmembrane helix 7 removed for clarity). Using identical camera coordinates, the silhouettes of these extremum pockets at the receptor origin (middle spline) are drawn to provide examples of different shapes and volumes across different genes and states.

Moreover, in five apo state experiments, no orthosteric binding site was detected by the automated pipeline. This occurs because the cavity detection algorithm applies strict spatial criteria - specifically requiring candidate pockets to maintain the majority of their volume in proximity to the reference ligand centroid. In these specific apo trajectories, the algorithm either identified cavities that were too small and shifted laterally away from the reference site or detected overly diffuse, oversized voids that extended deep into the core or out toward the extracellular milieu, thereby failing to meet the required spatial threshold.

Plotting how the volume changes across simulated frames (Figure 2 A) further highlights the need for MD: the binding site shows a very dynamic behaviour in both states and all receptors, which is in accordance with the RMSD values of the binding site residues discussed above.

The dynamic nature of identified cavities is evident in the highly heterogeneous volume classification: while the majority (66.4%) fall into the large or medium-sized categories, 13.3% are categorised as small, and 20.4% as very large (Figure 2 A). This wide range is partly influenced by methodology, as orthosteric cavities occasionally extend into the extracellular vestibule and absolute cavity volumes depend on the geometric definitions and algorithms used (mdpocket [13]). The volumes reported should therefore primarily be interpreted as relative measures obtained within a consistent computational framework, rather than as absolute physical cavity volumes. Nevertheless, because all trajectories were analysed using the same workflow, comparisons between cavities provide a consistent measure of relative changes in their dimensions.

The volume of the smallest holo orthosteric pocket (139 Å^3^) is almost double the volume of the smallest apo orthosteric pocket (82 Å^3^), even though their shape and 3D location are similar. Although the largest apo orthosteric pocket (1014 Å^3^) is notably very large, it is still smaller than the largest holo orthosteric pocket (1251 Å^3^) and has a different shape and 3D orientation. It seems as if the binding site residues have moved to close off at least part of the pocket to a funnel-like shape, whereas in the holo conformation, the binding site is fully open and available for the ligand to occupy (Figure 2 D). It must be noted that the largest five apo pockets at the orthosteric site are in fact larger than expected and close in volume to the holo orthosteric pockets. However, the shape of the pockets can vary drastically.

Together, the orthosteric binding site analysis clearly indicates the inherent plasticity of the pocket and the ligand-specific induced dynamics, reinforcing the necessity of MD-based approaches for accurate pocket characterisation.

## 3. TAAR Pocket Dynamics

Having established how receptor dynamics and ligand binding shape the orthosteric site, we explored whether similar conformational variability extends across the wider receptor surface by mapping the allosteric pocketome.

Beyond the well-characterised orthosteric binding site, GPCRs frequently utilise allosteric sites to modulate receptor signalling, offering a pathway for highly selective ligand design [20], [21]. Experimentally characterised non-orthosteric binding regions have been identified at diverse positions within GPCR structures, including extracellular vestibules, intracellular regions and membrane-facing sites, as well as lipid-associated cavities such as cholesterol-binding regions [22], [23], [24]. Unlike the orthosteric pocket, these allosteric cavities are often transient and may not be captured in static experimental snapshots, such as X-ray crystallography or cryo-EM [20]. Consequently, the receptor’s dynamic motion is a prerequisite for revealing or closing these sites over time.

Despite the growing importance of allosteric regulation, simulation-based characterisation of pocket dynamics across the TAAR family remains underexplored. Given the relatively recent emergence of structural data for this receptor family, there is a lack of a structured, comparative analysis between apo and holo states regarding ligand-induced conformational changes within the pocketome.

To address this gap, we analysed 2,364 identified cavities across our entire simulation library, providing a high-resolution, time-resolved view of the TAAR pocketome. Importantly, the cavities identified by the present geometric analysis should not automatically be interpreted as functional ligand-binding or allosteric sites. Cavity detection establishes the presence and dynamics of geometrically accessible regions, whereas their ability to accommodate a ligand and their functional relevance require further computational or experimental validation.

Our analysis reveals that approximately 27% of all identified cavities exhibit transient behaviour, defined here as an absence in at least 10% of simulation frames (100 ns) and a consecutive absence of at least 15 ns (Figure 3B). Notably, this transiency rate is independent of the ligand state, remaining consistent across both apo and holo simulations. We observed that transiency is largely driven by receptor-specific structural variation rather than gene-family categorisation. For instance, hTAAR1, mTAAR1, and mTAAR9 display remarkably uniform transiency rates (25%, 25%, 26%, respectively). A notable exception is the murine TAAR7f, which exhibits a higher transiency rate of 30.0% (apo) and 34.8% (holo), resulting in a mean of 33%. This variance is likely influenced by limited structural sampling, as this target is represented by a single PDB structure, yielding a smaller median pocket count (14 in apo, 17 in holo). Critically, however, transient behaviour is a feature observed in every replicate, confirming that this dynamic opening and closing of cavities is an intrinsic property of the TAAR fold.

**Figure 3.**
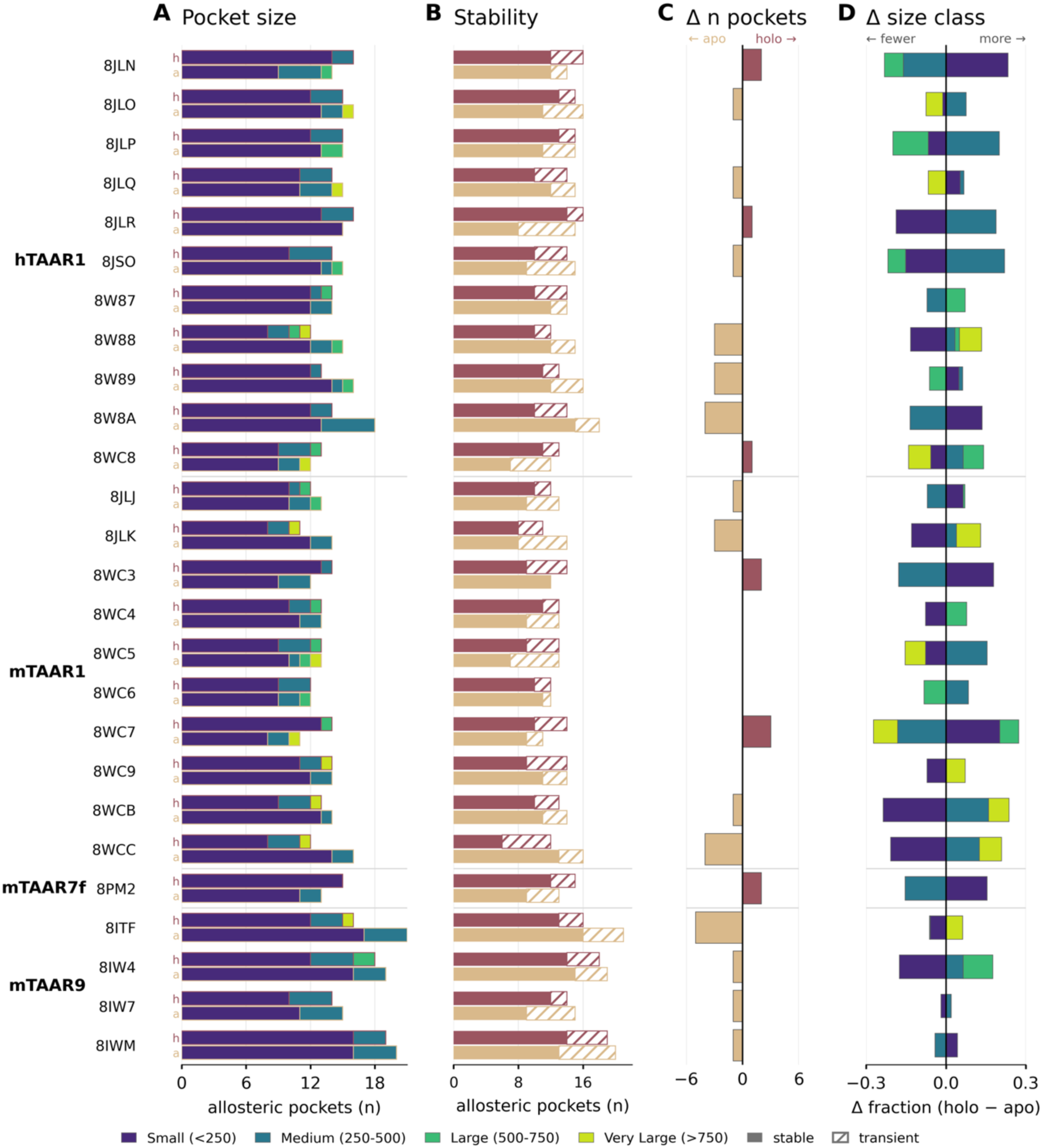
Characterisation of putative allosteric pockets. **(A)** Volume categories of all identified cavities. **(B)** Pocket transiency rates. **(C)** Difference in the number of pockets when comparing apo and holo states. **(D)** Difference in size categories (fraction of holo minus apo states) for all pockets that do not meet the orthosteric site classification.

Pocket transiency seems to correlate with cavity volume. Small pockets (defined as < 250 Å³) represent the majority of our dataset (∼ 75%) and exhibit a transiency rate of 33%. In contrast, larger cavities are significantly more stable: medium (250 – 500 Å³), large (500 – 750 Å³), and very large (> 750 Å³) pockets show transiency rates of 11%, 13%, and 15%, respectively (Figure 3).

While these distributions provide a global overview, the number and volume distributions (Figure 3A, 3D) of cavities vary even across individual experimental setups (PDB ID - replicate – state). A volume category differential plot (Figure 3D) illustrates how the pocketome is remodelled in the presence of the ligand. For example, the structure with PDB ID 8JLN has more small pockets and fewer medium and large cavities in the holo simulation than in the apo simulation. However, these trends across size categories do not show a single pattern but depend on the individual structure analysed. The holo state has more small cavities than the apo state in only 30% of structures. Similarly, 23% of structures have more large cavities detected in the holo state. Instead, looking at the medium-sized barrel, 58% of structures have more cavities of that size reported in the holo state.

Taken together, our data strongly suggest that the pocketome is a dynamic and ligand-dependent system of cavities spread across the receptor surface. This heterogeneity highlights the complexity of the TAAR pocketome and suggests that localised structural rearrangements - rather than global gene-family motifs - govern pocket availability.

## 4. TAAR Global Pocket Identification

Because pocket identifiers generated by individual mdpocket analyses are local to each run, identical numerical labels do not imply spatial correspondence across independent simulations. For example, pocket 07 in apo 8JLR replicate 1 does not necessarily occupy the same region as pocket 07 in apo 8JLR replicate 2. Thus, although each MD simulation provides a detailed description of its individual pocketome, the local numbering scheme does not establish pocket identity across simulations.

To enable direct comparison of cavities between independent simulations, we introduced a Global Identifier (GID) based on absolute 3D position relative to a shared, aligned receptor origin. Introducing a global ID would mean that identity is based on the 3D space in which the receptor and its pocketome reside, rather than a per-run label. Although the GID is conceptually better than a local label, it can only be used in a meaningful dataset. This limited dataset must use the same reference space, the receptors need to be aligned with one another, and, most importantly, if the subset is not chosen wisely, the GID will saturate until it eventually maps the entire surface area of the receptors relative to one another. The comparability we provide is achieved by selecting a meaningful subset of the data. To validate the GID approach, we established a targeted subset comprising 24 experiments across four genes: 8JLR for hTAAR1, 8JLJ for mTAAR1, 8PM2 for mTAAR7f, and 8ITF for mTAAR9, spanning both the apo and holo states with three replicates each

We first assessed whether the GID framework recovers a structurally defined reference region without prior information about its functional identity. Across the combined 24-simulation dataset, the orthosteric pocket was consistently assigned to GID3 across all replicates and structures (Figure 4). This is notable because the voxelised IoU graph-based method is agnostic to the orthosteric pocket location, and the orthosteric site frequently fragments into smaller sites in apo simulations. Its consistent assignment to a common global region, therefore, provides a positive control indicating that the GID approach can recover a conserved spatial cavity region from independently detected local pockets.

**Figure 4.**
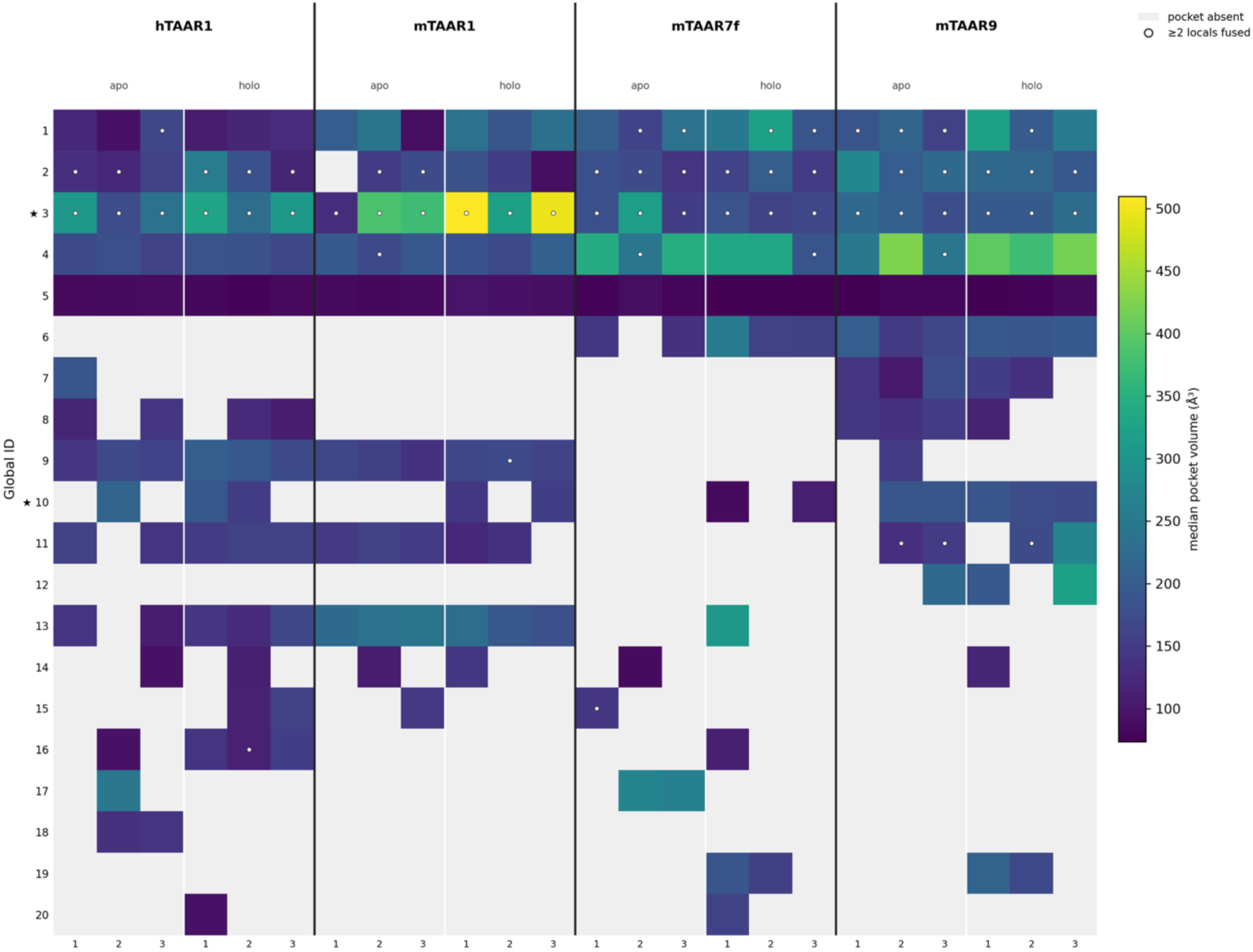
Heatmap of global ID presence across the 24-simulation dataset. The viridis colour scale is used to depict the summed volume of the local pockets contained by this GID. The stars signal GID members that are in touch with binding site residues.

We next tested whether similar global regions were recovered when the apo and holo datasets were processed independently. Both analyses returned 22 GIDs that map onto each other to a large extent; 21 of these had a corresponding centroid within 10 Å. Seventeen of these represented reciprocal nearest-neighbour matches, with a median centroid separation of 1.35 Å. It should be noted that only 4 of these 17 reciprocal matches share the same GID number - underscoring that GID numbers are arbitrarily assigned within the chosen dataset. Nevertheless, it is a promising result to have found the same global pocketome regions in independent runs. The non-reciprocal matches and non-match can also have a biological meaning, representing the difference between the apo and holo states. Furthermore, the pockets that have a matched GID between the individual runs sit approximately 5 Å from their GID-mates but 10 Å away from the nearest neighbouring GID.

Having established that the GID framework can recover comparable spatial regions in independently analysed datasets, we next used it to assess pocketome reproducibility across MD replicates (Figure 5). While the number of local pockets, i.e., the ones defined by mdpocket within an experiment, found per replicate differs quite drastically, the number of GIDs shows less variation (Figure 5A). This alone does not elucidate pocketome changes, though, as, e.g., pockets can separate from one another even if they remain within the same global area. When analysing the replicates in a single GID run, it becomes apparent that the number of GIDs remains within the range of the local pockets. Therefore, detecting more pockets in one replicate does not mean sampling a larger pocketome; rather, it reflects fragmentation of the detected cavities.

**Figure 5.**
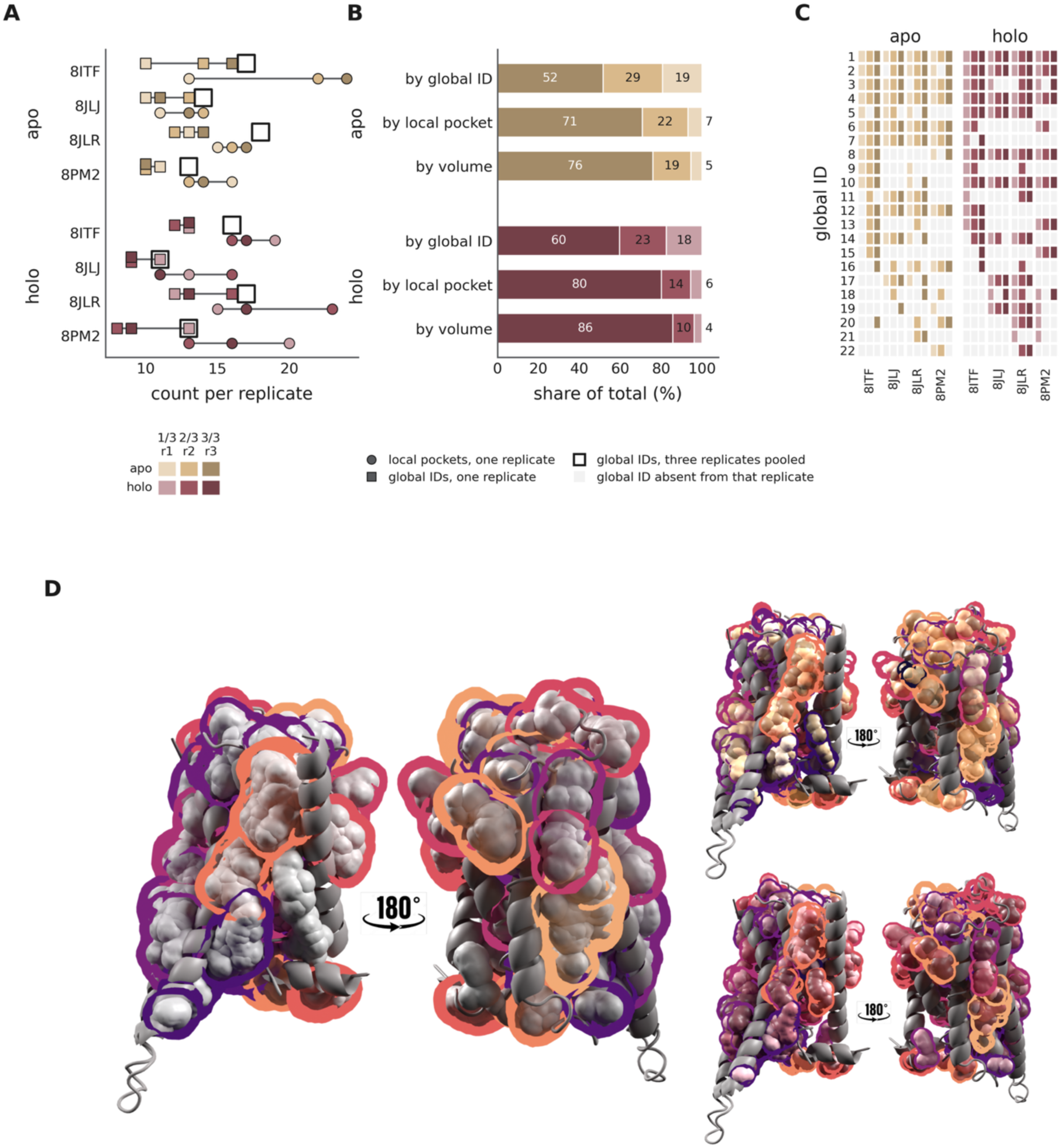
Analysis of replicate concordance of the reduced example cases in apo and holo state. **(A)** Comparison of the number of pockets found (circles) versus the mapped global identifiers (filled squares) and the total number of distinct GIDs found across all replicates. **(B)** Occupancy composition showing how many pockets are found in all three replicates (darker colour), two out of three replicates (medium colour) or one out of three replicates (lighter colour), categorised by their global ID, local ID and their volume. **(C)** Presence matrix mapping the GIDs to their presence in this receptor, replicate and state. **(D)** 3D context of GIDs mapped onto one exemplary receptor structure (8JLR). The coloured envelopes show which regions are summarised by GIDs. The left panel shows a combined view of the apo-only and holo-only runs, with pockets in grey and the matched GIDs as envelopes in shades of magma. GIDs found in only one of the two states (not matched) are not shown here. The two smaller receptor structures show the same information divided by their state, i.e., only the apo (sand, upper panel) or only the holo (mauve, lower panel) pockets with their replicates coloured by intensity and their GID as envelopes in shades of magma.

Classifying GIDs according to their occurrence across replicates further separated reproducibly sampled cavity regions from more variable regions (Figure 5C). Roughly 52% (apo) and 60% (holo) of GIDs are detected in all three replicates, 29% and 23%, respectively, are shared in 2/3 replicates and 19% and 18% are singleton GIDs. The contribution of these categories changed when the cavity volume associated with each GID was considered: 76% (86% in holo) of volume is explained by the GIDs that are shared in all replicates and only 5% (4% in holo) of the volume is associated with singleton GIDs. Therefore, the changes in the pocketome across replicates are driven by individual small pockets (Figure 5B).

The 3D representation of these GIDs further illustrates how multiple local cavities from independent simulations can occupy common global regions (Figure 5D). GIDs do divide the receptor’s pocketome into distinct global regions that can be shared across apo and holo local pockets of their three replicates. The left section is a combined view of only the matched GIDs (magma-sampled envelopes) and both apo and holo local pockets of all replicates (shown in grey). The right section shows the matched GIDs with only apo local pockets (top) or only holo local pockets (bottom). Since the envelopes stay the same, the right section shows how different fractions of the GID are occupied by apo versus holo pockets. Together, these observations show that raw cavity counts can overestimate differences between MD replicates because a shared spatial region may fragment into different numbers of locally detected cavities. Spatial grouping through the GID, therefore, reduces this complexity and provides a more informative representation of replicate concordance than local cavity counts alone.

It should be noted that singleton GIDs can also be a biological result – a true sampling of movement in the pocketome that, by chance, occurred in only 1/3 of replicates. The GID framework should therefore complement rather than replace expert structural analysis. Its purpose is to reduce large sets of trajectory-specific cavity assignments to comparable spatial regions, thereby facilitating the identification of reproducibly sampled regions, as well as potentially interesting replicate-specific events for subsequent analysis.

We finally applied the GID framework to compare cavity sampling between the apo and holo states within our 24-simulation dataset (Figure). Figure 6A highlights the identity of the global pockets by using envelopes around local pockets (shown in their state colours, sand (apo) and mauve (holo)) and giving the GID as text. While some pockets are clearly populated by both states, others remain state-dependent. Since the proposed GID is data-driven and valid only within its run, it provides the opportunity to compare the pocketome of a chosen dataset. By looking at the empty global envelopes in Figure 6B and 6C, respectively, the absence of a pocket becomes apparent.

**Figure 6.**
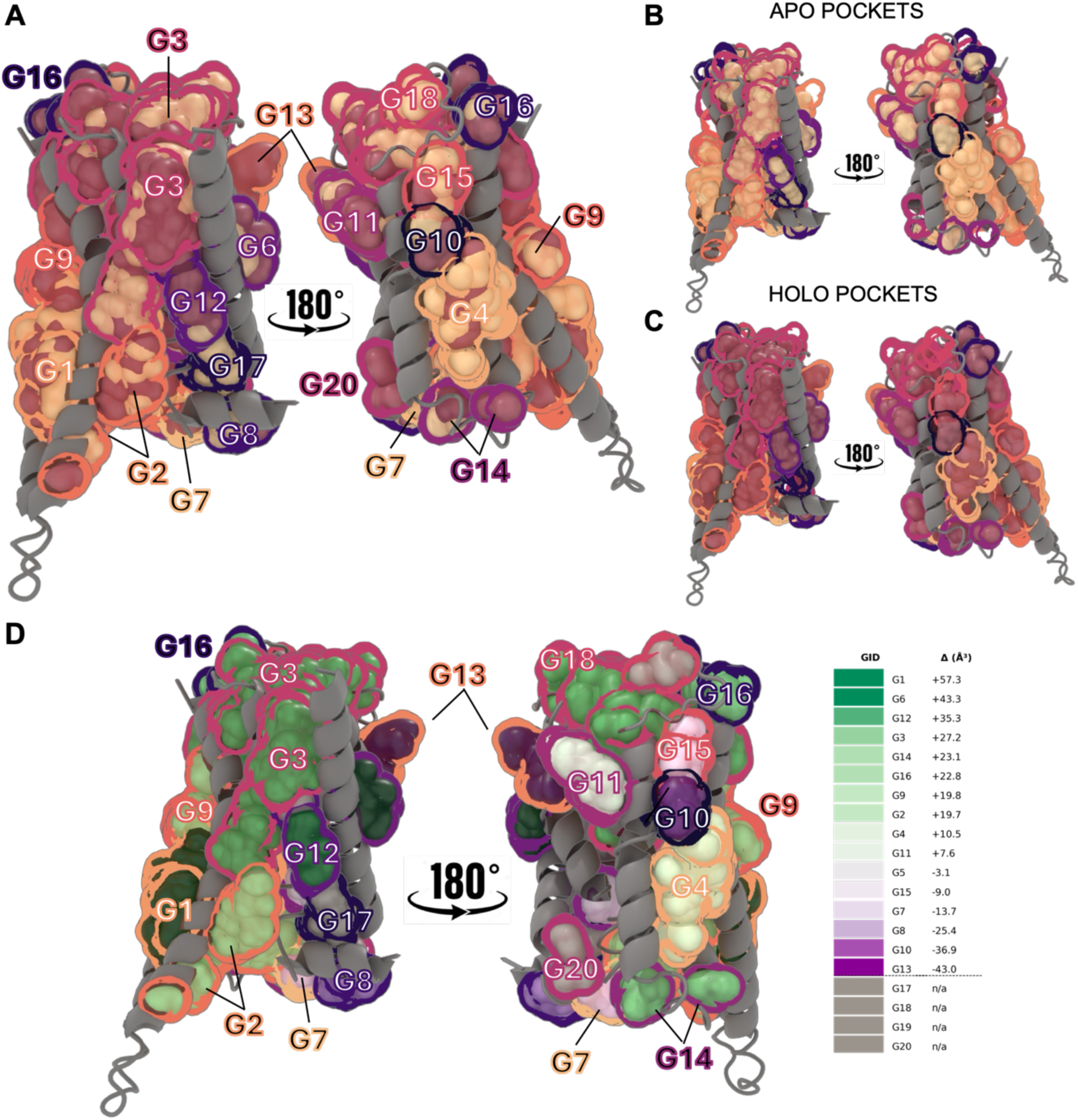
State-specific comparison of global pocket distribution between apo and holo trajectories. The GID is shown as envelopes around local pockets. Local pockets are coloured by their state (apo = sand, holo = mauve). **(A)** global IDs of both apo and holo state highlighted via text labels. Global pockets 5 and 20 are hidden by helices and other pockets, therefore not visible in this 2D projection. **(B, C)** GIDs in apo and holo states, respectively. The places in which the envelope appears to be empty are due to the absence of local pockets of the other state. They highlight in which places the apo and holo states differ more visibly. **(D)** Comparison of the volume changes between apo and holo states. The volume delta starts with a dark green colour for G1, meaning that this region has a larger volume in the holo state than in the apo state, and G13, which is larger in the apo state (shown by its purple colour). The envelopes around the global pockets are shown in the same colours as in Figure 4 and labelled. White-ish colours mean that the volume of this global region did not change noticeably. G17 (apo), G18 (apo), G19 (holo), and G20 (holo) exist in only one state and are therefore coloured grey.

Figure 6D compares cavity volumes within matched GIDs between the apo and holo simulations. While envelope colours stay linked to their GID, the pockets are shown in a green to purple gradient, depending on their volume changes. The GID is used to determine whether the local pockets within it have changed in size when the ligand is removed. Green pockets have a larger volume in their holo state, while purple pockets are larger in the apo state. Some global regions contain larger cavities in the holo state, others are larger in the apo state, while several exhibit comparatively little difference between states. Four GIDs were detected in only one of the two states and therefore could not be compared directly by volume. Overall, these observations indicate that removal of the co-bound ligand is associated with spatially heterogeneous changes in the TAAR pocketome rather than a uniform expansion or contraction of cavities across the receptor.

The GID framework does not establish a universal identity for individual cavities across TAARs or GPCRs more broadly. Instead, it provides a question-driven spatial identifier for comparing pocketomes within appropriately selected and structurally aligned datasets. In the present study, this enables two complementary applications: assessing which cavity regions are reproducibly sampled across independent MD replicates and identifying spatial regions that differ between ligand-bound and ligand-depleted simulations. By consolidating fragmented local cavity assignments into shared three-dimensional regions, the approach reduces the complexity of trajectory-level pocket data and facilitates targeted analysis of dynamic receptor cavities. Although developed and evaluated here using TAARs, the framework is not inherently TAAR-specific and can be applied to other structurally aligned receptor datasets for which dynamic cavities need to be compared across simulations.

## Conclusion

In this study, TAAR receptor plasticity was explored across 156 µs of MD simulations utilising 26 experimental structures from four human and murine TAAR genes, simulated in both apo and holo states. The resulting dataset provides a systematic view of TAAR structural dynamics, ranging from movement of the receptor and orthosteric binding site to changes across the broader receptor pocketome.

Analysis of the orthosteric binding site revealed structural plasticity and a state-dependent difference in cavity dimension. In most systems, the orthosteric cavity was larger in the holo state, whereas ligand removal frequently resulted in contraction or fragmentation of the binding site into smaller cavities. At the same time, binding site RMSD did not show a uniform apo–holo trend, indicating that ligand presence does not simply restrict local residue mobility. Together, these observations highlight a dynamic relationship between ligand occupancy, binding site geometry, and local conformational sampling. Extending the analysis beyond the orthosteric site revealed a heterogeneous landscape of non-orthosteric cavities distributed across the receptor structure, ranging from small and transient to larger and more persistent regions. Their occurrence and dimensions varied between simulations and states, highlighting the dynamic TAAR pocketome. Systematic identification of these cavities provides a structural framework for prioritising and exploring previously uncharacterised regions of TAARs and investigating their potential ligandability and functional relevance. Notably, transient pockets, which can only be seen and analysed in MD simulations, are reported in each experiment and in both states, providing a promising research target.

To enable systematic comparison of this dynamic cavity landscape across independent simulations, we introduced the GID framework, which groups trajectory-specific cavities according to their spatial identity within a selected, structurally aligned dataset. The GID analysis revealed that variability in raw cavity counts can overestimate differences between independent MD replicates, as shared spatial regions can fragment into different numbers of locally detected cavities. Most of the sampled cavity volume was associated with GIDs that were reproducibly detected across all three replicates, whereas replicate-specific GIDs accounted for only a small fraction of the overall volume. Independent GID assignments of the apo and holo datasets recovered a largely conserved set of spatial regions while retaining state-specific differences in cavity sampling, enabling differences in cavity occurrence and geometry to be analysed within a common three-dimensional framework. Thus, spatial identification provides information that cannot be obtained from trajectory-specific cavity labels or raw cavity counts alone and facilitates the identification and prioritisation of specific dynamic cavity regions for further investigation. Rather than providing a universal cavity nomenclature, the GID is designed as a question-driven approach for comparing appropriately selected and structurally aligned datasets.

Together with their spatial and dynamic characterisation, the resulting cavity maps provide starting points for future computational and experimental studies of potential non-orthosteric modulation. Beyond the biological findings presented here, the study establishes a computational resource for the broader investigation of the understudied TAAR family. The MD simulation library, three-dimensional maps of dynamic TAAR cavities, and accompanying computational analysis pipeline provide reusable resources for investigating receptor dynamics, ligand recognition, and structurally informed questions such as TAAR ligand discovery. Making the data and workflow publicly available will also enable researchers to reproduce, extend, and apply these analyses to new research questions. Although developed and evaluated using TAARs, the underlying approach is not inherently TAAR-specific and may be transferable to other structurally aligned receptor datasets, including other GPCR families. Together, the dataset and analytical framework establish a foundation for comparative, data-driven analysis of dynamic receptor pocketomes.

## Material and Methods

### System preparation and molecular dynamics simulations

The structures of human TAAR1 (11), murine TAAR1 (10), murine TAAR9 (4) and murine TAAR7f (1) were downloaded from the Protein Data Bank [25] (PDB, https://www.rcsb.org/) in August 2024.

An extensive list of the receptors, their ligands and the references can be found in Table 1. System preparation was performed in Maestro using the Protein Preparation Wizard [31] (Schrödinger Suite, release 2024-1) to remove potential nanobodies, G proteins, additional co-crystallised molecules and ensuring optimization of hydrogen bonds and side chains at physiological pH (pH = 7.4). Additionally, in case of missing substructures, homology modelling based on an AlphaFold multimer model was performed for gaps that especially exist in the intracellular loop 3. In apo simulations, the ligand was removed from the prepared holo structures using visual molecular dynamics (VMD) [32].

**Table 1.** human and murine TAAR structures used in this study, sorted by their gene with their PDB ID, co-bound ligand and primary publication listed.

| PDB ID | Gene name | Ligand | Ref. | PDB ID | Gene name | Ligand | Ref. |
| --- | --- | --- | --- | --- | --- | --- | --- |
| 8JLN | hTAAR1 | T1AM | [26] | 8WC3 | mTAAR1 | SEP363856 | [27] |
| 8JLO | hTAAR1 | Ulotaront | [26] | 8WC4 | mTAAR1 | ZH8651 | [28] |
| 8JLP | hTAAR1 | Ralmitaront | [26] | 8WC5 | mTAAR1 | Tetramethylammoniumion (TMA) | [28] |
| 8JLQ | hTAAR1 | fenoldopam | [26] | 8WC6 | mTAAR1 | PEA | [28] |
| 8JLR | hTAAR1 | A77636 | [26] | 8WC7 | mTAAR1 | ZH8667 | [28] |
| 8JSO | hTAAR1 | AMPH | [26] | 8WCC | mTAAR1 | Cyclohexylammoniumion (CHA) | [28] |
| 8W87 | hTAAR1 | Methamphetamine | [28] | 8WC9 | mTAAR1 | ZH8651 | [28] |
| 8W88 | hTAAR1 | SEP363856 | [28] | 8WCB | mTAAR1 | CHA | [28] |
| 8W89 | hTAAR1 | 2-Phenylethylamine (PEA) | [28] | 8PM2 | mTAAR7f | DMCH | [29] |
| 8W8A | hTAAR1 | RO5256390 | [28] | 8ITF | mTAAR9 | DMCH | [30] |
| 8WC8 | hTAAR1 | ZH8651 | [27] | 8IW4 | mTAAR9 | Spermidine (SPE) | [30] |
| 8JLJ | mTAAR1 | T1AM | [26] | 8IW7 | mTAAR9 | PEA | [30] |
| 8JLK | mTAAR1 | Ulotaront | [26] | 8IWM | mTAAR9 | PEA | [30] |

The orientation of TAARs in their POPC membrane was determined by a reference structure (8IW7, mTAAR9) downloaded from Orientations of Proteins in Membranes [33] (https://opm.phar.umich.edu/). The aligned receptors were then embedded into a pre-built 90 Å × 90 Å (with VMD Membrane Builder plugin 1.1) 1-palmitoyl-2oleyl-sn-glycerol-3-phospho-choline (POPC) square bilayer through an insertion method by using HTMD [34] (Acellera, version 2.2.7). Any lipids overlapping with the structure were removed and TIP3P water models were inserted. To keep the system at a physiological pH of 7.4 (concentration of 0.154 M), Na^+^ and Cl^-^ ions were added using VMD [32]. Simulations were performed with CHARMM36 force field [35]. A pipeline based on OpenMM (version 8.1) [36] utilising periodic boundary conditions in an NPT ensemble, i.e. with constant pressure and temperature during equilibration and a canonical NVT ensemble during production (once equilibrated) was employed. The settings for equilibration can be found in Table 2 and for the production in Table 3. A Langevin integrator was used for all simulations. In both the equilibration and production phases of MD, three replicates were run per PDB ID and state (apo/holo) combination, resulting in a total aggregate time of 156 microseconds.

**Table 2.**
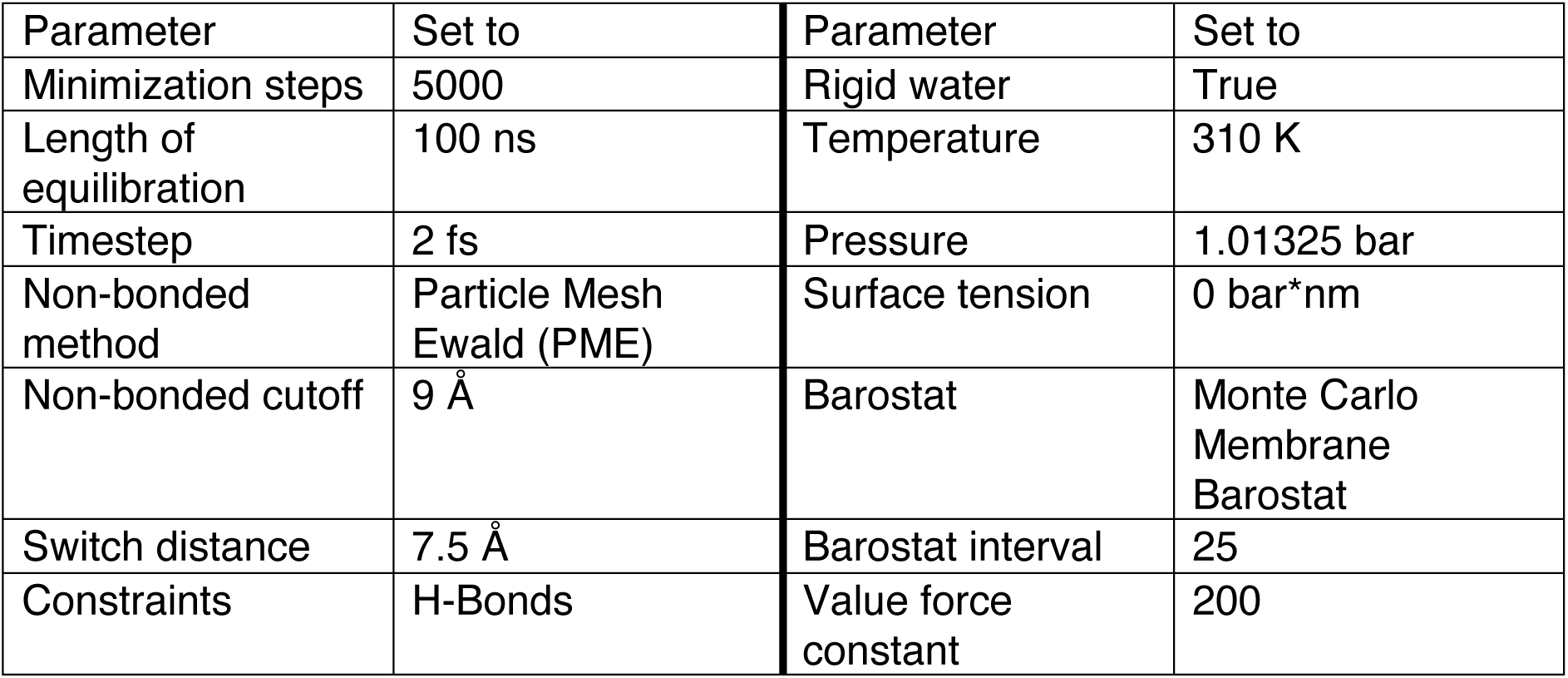
Equilibration settings for all MD simulations performed, giving the parameter name and its value.

**Table 3.** Production settings of all MD simulations performed giving the parameter name and its value.

| Parameter | Set to | Parameter | Set to |
| --- | --- | --- | --- |
| Length of production | 1000 ns | Surface tension | 0 bar*nm |
| Timestep | 4 fs (apo), 2 fs (holo) | Temperature | 310 K |
| Trajectory frequency | 25000 | Pressure | 1.01325 bar |
| Non-bonded method | Particle Mesh Ewald (PME) | Friction | 0.1 ps |
| Non-bonded cutoff | 9 Å | Switch distance | 7.5 Å |

### MD Analysis and Meta-Analysis

A pipeline was generated and employed to launch the simulations and analyse the trajectories based on OpenMM [36], MDanalysis [37] and mdpocket [38]. The pipeline is accessible via (https://github.com/DiPizio-Lab/dynamic_pocketome_TAARs). Root Mean Square Deviation (RMSD) of the backbone Carbon-alpha (Cα) as well as Root Mean Square Fluctuation (RMSF, not shown but reproducible using the separately published code) were calculated after drying and aligning the receptors to the corresponding frame 0 (the experimental structure). The calculations excluding ICL3 were derived from the gene-specific ICL3 residue numbers, ± 3 residues. To automate pocket detection, separation and description, a custom Python script was generated. Given the topology and trajectory, a first mdpocket run was performed. To be able to perform the second run that characterises individual cavities, the results of the first run had to be automatically separated. This was accomplished by a new <u>separation</u> <u>algorithm</u> based on finite-element mesh topology. Rod elements consisting of two nodes were generated to connect individual data points derived from the alpha spheres that mdpocket uses to define cavities in the topology across its trajectory. A rod element is generated if two data points are closer than a pre-defined distance parameter of 1.4 Å. In an iterative approach, all elements are inspected and both element nodes are assigned the lower node number of the two nodes [39]. Repeating this process until a full inspection of all elements yields no further change identifies the groups that are not connected to one another and hence, the cavities. After this separation, the second run of mdpocket, defining characteristics as described by Schmidtke et al. [38], was run, providing useful information for individual pocket data analysis. Alternatively, the user can decide to employ DBSCAN clustering implemented using scikit-learn [40], [41].

All pocket descriptors and coordinates are fed into the <u>meta-analysis pipeline,</u> which evaluates and compares the overall characteristics of the identified pockets and user-defined subsets. In this study, the Python-implemented pipeline was used to identify the orthosteric binding site pocket(s), describe transiency and volume trends and visualise the overlap in subsets of pockets, e.g., gene specificity or overlap between replicates of the same experiment, as well as the comparisons between the two states. In this study, the <u>orthosteric site</u> is defined as a cavity or cavities whose centroid (mean x/y/z position of all its dummy-atom coordinates, averaged over every frame of the trajectory) lies within a 5 Å radius of the co-resolved ligand centroid. In apo simulations, the centroid of the corresponding holo ligand structure is used to define the binding site. A cavity is deemed <u>transient</u> if its volume stays at 0 Å^3^ for a minimum of 100 ns and at least 15 consecutive ns (once within the simulation). Volume categories are simply defined by chunking the data into barrels of < 250 Å^3^, 250 – 500 Å^3^, 500 – 750 Å^3^ and everything above 750 Å^3^, for the categories of small, medium-sized, large and very large, respectively.

Additionally, the <u>global 3D identifier</u> (GID) was implemented: first, for each individual pocket, the coordinates of the alpha spheres defined by mdpocket are voxelised onto a shared grid (1 Å voxels). Every pair of pockets gets an Intersection over Union (IoU) score on their voxel sets. Then a graph is built with one node per local pocket and an edge whenever the IoU is greater than 0.3. Each connected component is assigned a single global ID. A certain amount of dilation and translation in 3D is allowed in this process for each pocket (dilation radius of 5 Å) to allow for minor positional shifts. Very small and or noisy graph connections are re-assigned based on a k-nearest neighbour approach (calculated using the centroid of each body). Each graph produced by this process will represent a unique cavity in the receptor’s topology. The local pocket IDs (defined by mdpocket and ATClus) are then extended by a new global ID, which enumerates the individual graphs. By this approach, the dataset will have both local information, i.e., separate cavity entities only valid in this PDB + replicate + state and global information based on the user-provided subset of experiments. To demonstrate the global ID in this study, one representative structure per gene is chosen, resulting in the subset of 3 replicates and two states of 8JLR (hTAAR1), 8JLJ (mTAAR1), 8PM2 (mTAAR7f) and 8ITF (mTAAR9), equalling 24 experiments. Notably, increasing the number of experiments considered increases the entropy of the dataset of alpha spheres and converges to model the surface area of the receptor with all its indents. Therefore, it is recommended to choose meaningful subsets, e.g., the replicates of the same structure and state, to analyse the statistical sampling of movement in MD simulations or the apo and holo state of the same structure to compare structural differences induced by ligand presence.

#### Replicate analysis

In three separate runs, four representative PDB structures (8JLR, 8JLJ, 8PM2, 8ITF) in two states, and three replicates of each, are used to evaluate replicate concordance. GIDs are only valid within their run, i.e., within the apo-only, holo-only, or both states run. Per structure and identifier, the occurrence of replicates is counted (3/3 = core, 2/3 = shell, 1/3 = singletons) and expressed as identifier counts, local pocket counts and summed interpolated median volume. Apo and holo only GIDs are matched on member-pocket centroids by finding the pair with the smallest Euclidean distance to each other. The Euclidean distance of centroids was built in both directions and resulted in reciprocal matches, unidirectional matches or non-matches if the centroids were separated by more than 10 Å.

#### GID delta volume

The algorithm pulls every local pocket associated with the given global ID (GID), split by state. Then, the median of all pockets’ median interpolated pocket volume is calculated and compared across states. Therefore, the presented value is calculated as the median of the differences between local holo and apo pocket volumes.

The meta-analysis scripts will generate a database-like summary of all pockets, their coordinates across the number of frames and their descriptors in addition to the local and global ID, the newly calculated transiency and volume category. Furthermore, new flags indicate if the pocket is near the ligand orthosteric binding site, the largest pocket in this experiment and how many frames remain continuously at zero. The visualisations shown in this paper are also generated by the Python workflow utilising matplotlib [42] and plotly [43] libraries. All 3D renders were performed in Blender [44] using the molecular nodes [45] extension.

#### Use of AI

Claude Code [46] was used in this paper to find redundant code fragments and help write precise docstrings in addition to minor code edits. Grammarly [47] was used to spell- and grammar-check the text of this paper. Additionally, for minor edits and text corrections, Microsoft Copilot was used [48].

