## Supplemental Table and Figure for "Dynamic Pocketome of Trace Amine-Associated Receptors"

**Supplementary Data**

Sup Table 1 Comparison of apo and holo experiments in terms of RMSD. The table lists the difference of RMSD CA and RMSD of the side chains of the binding site with the apo values as references.

| PDB ID | Gene | Diff. RMSD protein Cα | In % | What moves more | Diff. RMSD binding site side chains | In % | What moves more |
|---|---|---|---|---|---|---|---|
| 8ITF | mTAAR9 | -0.169 | -5.77% | apo | -0.589 | -14.56% | apo |
| 8IW4 | mTAAR9 | 0.3175 | 8.45% | holo | -0.446 | -9.12% | apo |
| 8IW7 | mTAAR9 | 0.31 | 10.01% | holo | 0.3565 | 9.78% | holo |
| 8IWM | mTAAR9 | -0.4215 | -13.85% | apo | 0.07 | 1.78% | holo |
| 8JLJ | mTAAR1 | -0.06 | -2.48% | apo | 0.0605 | 3.29% | holo |
| 8JLK | mTAAR1 | 0.197 | 7.57% | holo | -0.5755 | -27.57% | apo |
| 8JLN | hTAAR1 | 0.0825 | 3.33% | holo | -0.1805 | -7.71% | apo |
| 8JLO | hTAAR1 | 0.198 | 7.06% | holo | 0.344 | 11.03% | holo |
| 8JLP | hTAAR1 | 0.393 | 11.56% | holo | 0.45 | 19.94% | holo |
| 8JLQ | hTAAR1 | 0.168 | 5.09% | holo | 0.1165 | 4.21% | holo |
| 8JLR | hTAAR1 | 0.4655 | 14.75% | holo | 0.462 | 18.66% | holo |
| 8JSO | hTAAR1 | 0.239 | 8.56% | holo | -0.424 | -14.49% | apo |
| 8PM2 | mTAAR7f | 0.7255 | 24.29% | holo | 0.0075 | 0.18% | holo |
| 8W87 | hTAAR1 | 0.6635 | 20.76% | holo | -0.1585 | -7.00% | apo |
| 8W88 | hTAAR1 | 0.4315 | 13.53% | holo | -0.545 | -22.29% | apo |
| 8W89 | hTAAR1 | -0.539 | -18.09% | apo | -0.301 | -12.31% | apo |
| 8W8A | hTAAR1 | -0.0525 | -2.17% | apo | -0.287 | -12.39% | apo |
| 8WC3 | mTAAR1 | -0.52 | -20.80% | apo | -0.099 | -5.11% | apo |
| 8WC4 | mTAAR1 | 0.981 | 29.40% | holo | -0.564 | -25.97% | apo |
| 8WC5 | mTAAR1 | 1.0765 | 29.65% | holo | -1.1775 | -37.46% | apo |
| 8WC6 | mTAAR1 | -0.0365 | -1.59% | apo | -0.0655 | -3.64% | apo |
| 8WC7 | mTAAR1 | 1.391 | 37.25% | holo | -0.6725 | -26.94% | apo |
| 8WC8 | hTAAR1 | 0.094 | 3.66% | holo | 0.126 | 6.57% | holo |
| 8WC9 | mTAAR1 | 0.1015 | 2.69% | holo | 0.2965 | 15.64% | holo |
| 8WCB | mTAAR1 | 0.245 | 9.36% | holo | 0.1 | 5.45% | holo |
| 8WCC | mTAAR1 | 0.1885 | 6.87% | holo | -0.4995 | -20.07% | apo |

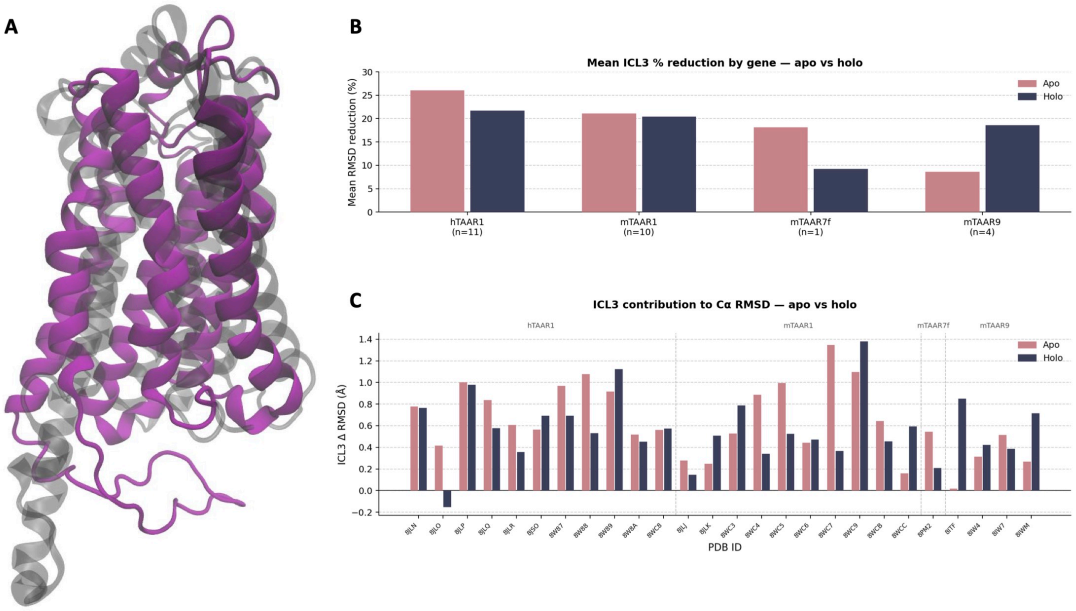


Sup Fig 1(A) highlights the movement of ICL3; shown is the maximal deviation of any experiment (8WC9, replicate 3) where the grey structure is the starting frame and the purple structure is the frame with the highest RMSD. (B) and (C) highlight the influence of ICL3 on the overall RMSD of CA. (B) shows a comparison of reduction in percent between apo and holo experiments, while (C) gives details on each experiment run and its RMSD reduction in Ångström.
